# Functional ultrasound imaging of the spreading activity following optogenetic stimulation of the rat visual cortex

**DOI:** 10.1101/2021.02.05.429985

**Authors:** M. Provansal, G. Labernede, C. Joffrois, A. Rizkallah, R. Goulet, M. Valet, W. Deschamps, U. Ferrari, A. Chaffiol, D. Dalkara, J.A. Sahel, M. Tanter, S. Picaud, G. Gauvain, F. Arcizet

**Author notes:** Equal contribution (G. Gauvain, F. Arcizet).

## Abstract

Optogenetic stimulation of the primary visual cortex (V1) is a promising therapy for sight restoration, but it remains unclear what total cerebral volume is activated after surface stimulation. In this study, we expressed the red-shifted opsin ChrimsonR in excitatory neurons within V1 in rats, and used the fine spatial resolution provided by functional ultrasound imaging (fUS) over the whole depth of the brain to investigate the brain response to focal surface stimulation. We observed optogenetic activation of a high proportion of the volume of V1. Extracellular recordings confirmed the neuronal origin of this activation. Moreover, neuronal responses were even located in deep layers under conditions of low irradiance, spreading to the LGN and V2, consistent with a normal visual information process. This study paves the way for the use of optogenetics for cortical therapies, and highlights the value of coupling fUS with optogenetics.

## Introduction

Optogenetics has revolutionized investigation of the central nervous system^1^, providing hope for the treatment of a number of conditions, including deafness^2,3^ or vision loss^4^. Optogenetic therapy is already widely applied to retinal cells to restore vision for *in vivo* light application^5– 11^, and is currently being assessed in two different clinical trials^12,13^. However, different approaches, targeting cells other than those of the retina, are required for diseases causing degeneration of the optic nerve (e.g. glaucoma) and for advanced retinal degeneration (late AMD). For such conditions, direct stimulation of the primary visual cortex (V1) is a promising alternative strategy. Indeed, high performance rates have been reported for the detection of forms, with great accuracy, by blind human patients with cortical implants^14^, and cortical electrical prostheses have been shown to elicit visual percepts and to alter visual behavior in nonhuman primates (NHP)^15^. Successes have been achieved with implantable devices, but this approach nevertheless has a number of serious drawbacks: invasive surgery, signal degradation over time, and a lack of cell-type specificity. In this respect, optogenetic therapy stimulating V1 at the cortical surface could potentially afford similar benefits, but with less invasive administration, stable expression over time and precise genetic targeting of the appropriate cell population. Two of the key aspects of this strategy are the activation of cortical layer IV neurons, as these are the first cells to receive visual information from the visual thalamus^16^ and the propagation of activity to other visual structures, which would favor the generation of visual percepts.

Layer IV is located deep in the cortex (>1 mm in NHP). It is therefore a major challenge to read and write neuronal activity, to demonstrate the efficacy of stimulation. Red-shifted opsins are very good tools for neuronal stimulation, as they make it possible to use lower light power to activate deeper neurons than blue-sensitive opsins, whilst also making it possible to use higher light intensity safely^17–19^. Electrophysiological recordings can report neuronal activity with unmatched spatiotemporal resolution, but over a very small spatial area^20^. Conversely, techniques such as optical imaging and fMRI have been coupled with optogenetics to report activity throughout the brain, but at the expense of a loss of both spatial and temporal resolution^21,22^. Like fMRI, ultrafast functional ultrasound imaging (fUS) can provide brain-sized maps of neurovascular activity changes, with a high spatiotemporal resolution (100 µm x 100 µm, 1Hz) even in deep structures (up to 1.5 cm)^23^. This technique has been used to investigate sensory processing in anesthetized^24^, awake^24–26^ and asleep^27^ rodents, and in behaving primates^28,29^. fUS imaging and electrophysiological recordings can, therefore, be used to describe the dynamics of local neuronal activity accurately whilst scanning neurovascular activity over the entire brain. We used these two techniques to determine whether the optogenetic stimulation of V1 at the cortical surface could induce neuronal activity even in deep cortical layers, and initiate the propagation of information to other visual structures.

We show here that red light stimulation at the cortical surface can activate visual neurons localized in deep cortical layers without triggering a thermal hemodynamic response and toxicity. We were also able to follow the propagation of this information in other visual brain structures (i.e. LGN and V2). More generally, this work shows that fUS imaging has the potential to provide a clear, brain-wide mesoscopic view of the neuronal activities resulting from optogenetic stimulation.

## Results

### fUS imaging of V1 optogenetic activation in rats

We used the optogenetic actuator ChrimsonR for visual restoration in the cortex, because of its red-shifted opsin properties^17^ and because it was already being used for visual restoration in the primate retina^11^ and had given promising results in clinical trials^13^. We maximized the optogenetic activation of V1 by using the CaMKII promoter to ensure expression limited to the excitatory neurons of V1. Indeed, a ubiquitous promoter might lead to the silencing of pyramidal neurons through the recruitment of inhibitory neurons. ChrimsonR was fused to the fluorescent reporter tdTomato to facilitate the visualization of transfected areas. Following preliminary screening, we used the AAV9-7m8 mutated viral capsid to express ChrimsonR in the V1 neurons of Long-Evans rats (Fig. 1a). The mean rate of neuronal transfection was 5.5% over all cortical layers (Fig. S1). ChrimsonR expression was not restricted to the soma, but spread to the axons and dendrites (Fig. 1a). We assessed the efficacy of the optogenetic stimulation of V1 cortical neurons, by using fUS imaging to measure brain activity in a large proportion of the brain: from AP −3.5 mm to AP −8 mm, the zone in which most of the early visual system areas are located. Activity in the cortical layers was assessed following either direct stimulation of the contralateral eye with a white LED (58 mW.cm^−2^), or stimulation at 595 nm delivered with an optic fiber placed at the surface of the transfected or non-transfected V1 areas (7 mW, ∼140 mW.mm^−2^ at the brain surface) (Fig. 1b). We chose to use durations and magnitudes of parameters similar to those previously used^24,30^ for stimulation of the eye or cortex (2 s at 4 Hz or 20 Hz for stimulation of the eye and cortex, respectively, separated by a 13 s period of darkness. This cycle was performed 20 times). For eye stimulation (Fig. 1b, left), we first imaged V1 with a single imaging plane at AP −7.5 mm and constructed an activation map including only pixels displaying significant CBV (cerebral blood volume) responses (*p*<0.05 with Bonferroni-Holm correction, Wilcoxon signed-rank test, one-tailed, relative to baseline activity). We detected strong activation in both the ipsilateral and contralateral superior colliculi (SC) (ipsilateral, *n*=121/488 activated pixels; contralateral, *n*=134/421 activated pixels), but almost no activation in the ipsilateral and contralateral V1 areas (ipsilateral, *n*=8/634 activated pixels; contralateral, *n*=9/370 activated pixels). The lack of response in both V1 areas and the strong signal in both SC may reflect the retinotectal nature of most rodent retinal outputs^16^ or an effect of anesthesia^31^. An increase in CBV was already clearly visible on single-cycle responses (gray dashed lines), as illustrated for the significant pixel (#14-92) in the ipsilateral V1 area (insert). Following direct stimulation of the contralateral eye, the mean response (black curve) peaked 2 seconds after the two-second stimulation represented by the patch in gray (mean: 19.8 ± 18.3%). For cortical stimulation (Fig. 1b, center), we observed a broad activation of the ipsilateral V1 area (*n*=310/634 activated pixels). The activation spread out of the V1 area at each border (medial and ventral) with V1 projections onto other visual areas. Single-cycle responses of the same example pixel showed larger, less variable increases in CBV variation than for stimulation of the contralateral eye (optogenetic, mean: 65.8 ± 32.6%, *p*<0.0001, Mann-Whitney). A previous study showed that blue light delivery to the brain could itself generate non-specific changes in CBV ^30^. We therefore performed a control stimulation by locating the optic fiber at the surface of the non-injected V1 area, which did not express ChrimsonR-tdT (Fig. 1b, right). We used the same light stimulation parameters for this control as before. We detected no significant CBV responses in the non-injected V1 area under such optogenetic stimulation conditions.

**Figure 1.**
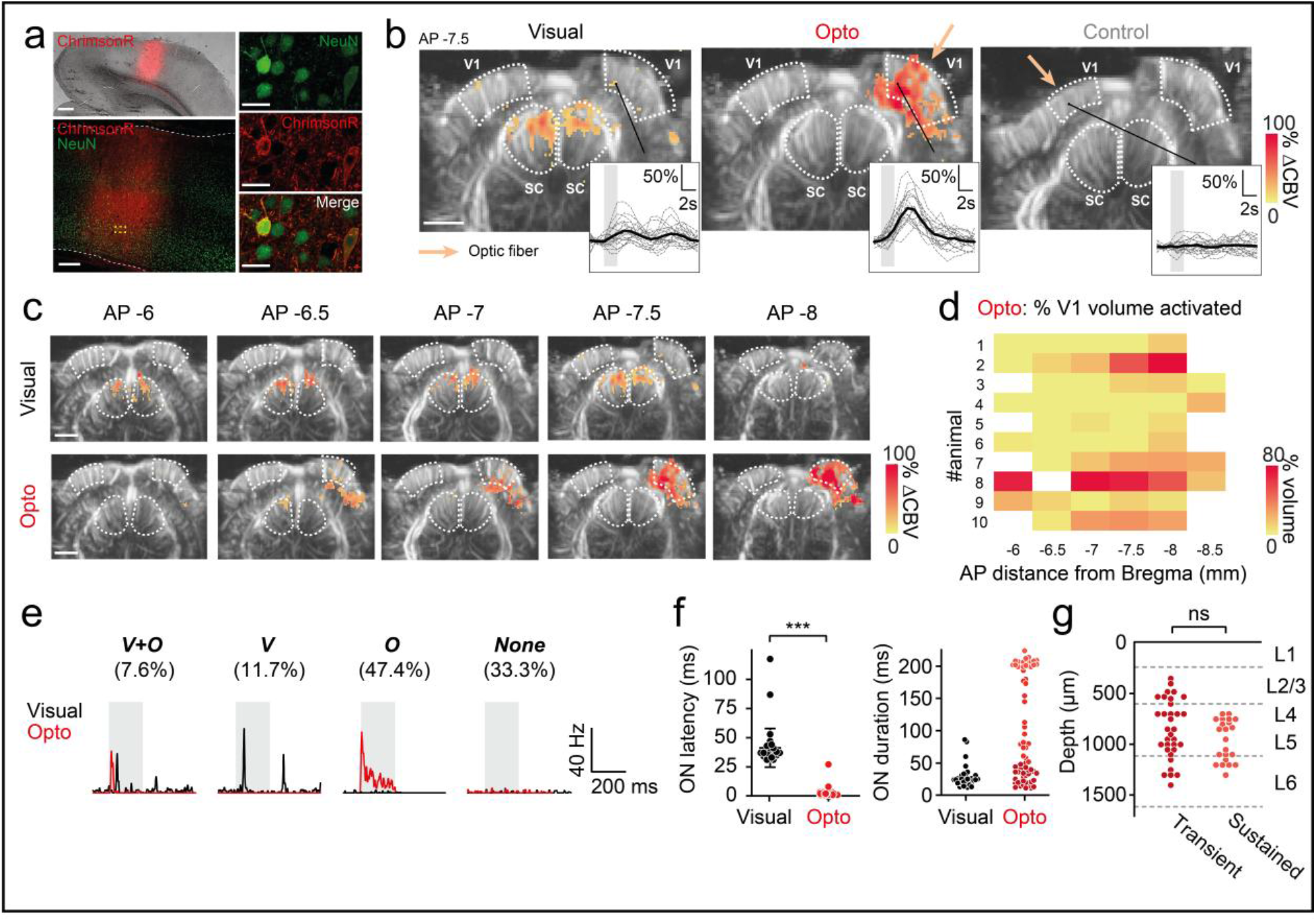
V1 neurovascular and neuronal activations resulting from cortical surface stimulation in rats. (a) Left: 2.5x imaging (top) and 20x confocal imaging of a brain section showing the localization of ChrimsonR in V1. White dashed lines delimit the cortical surface and border between V1 and the white matter. Right: close-up (40x) of the area delimited by the yellow lines in the previous image, showing two ChrimsonR-expressing neurons. Scale bars: 500 µm, 200 µm, 20 µm. (b) fUS activation maps obtained after visual stimulation of the contralateral eye (left), optogenetic stimulation of the ipsilateral V1 area expressing ChrimsonR (middle) and control optogenetic stimulation of the uninjected contralateral V1 area (right), from a single imaging plane (AP −7.5 mm) from the same animal. White dashed lines delimit the V1 and SC areas. Colored pixels indicate significant CBV variation (Wilcoxon signed-rank test with Bonferroni-Holm correction). Right insets: patterns of single-pixel activation. Gray lines represent single-trial activity (*n*=20) and the black line represents the mean CBV variation. Colored patches indicate light stimulation (duration: 2 s) (c) fUS activation maps for visual and optogenetic activation, for all imaging planes (AP −6 mm to AP −8 mm) in which V1 (delimited by white dashed lines) is present, from the same animal (animal #2). (d) Left: percentage of V1 activated during optogenetic stimulation for each fUS imaging plane, for all animals (*n*=10). Right: AP distribution of the ChrimsonR expression area on brain sections. (e) Spike density function (SDF) of typical V1 single units during visual (black lines) or optogenetic (red lines) activation. Four subpopulations of neurons were identified: double-responsive (n_V+O_=13), responsive to the visual (n_V_=20) or optogenetic stimulus only (n_O_=81), non-responsive (n_None_=57). (f) Left: ON response latencies for visual (*n*=33 units) and optogenetic (*n*=76 units) stimulation (Mann-Whitney, *p*<0.0001). Right: ON response durations for visual (*n*=33 units) and optogenetic stimulation (*n*=72 units). (g) Depth profile of transient (ON duration<51 ms, *n*=33 units) and sustained (ON duration>197 ms, *n*=22 units) neurons activated by optogenetics. (c-d) Scale bar: 2 mm. The irradiance used for optogenetic and control stimulation was 140 mW.mm^−2^.

We further characterized the V1 activation volume generated by contralateral eye stimulation or by stimulation of the transfected V1 area, by imaging all the planes containing V1 (from AP −6 to −8 mm, Fig. 1c-d). As shown in Fig. 1b, direct eye stimulation induced CBV responses mostly in the SC areas, but very little activation was observed in the two V1 areas (mean: 0.05 ± 0.12%). Strikingly for the same animal shown (as shown in Fig. 1c), direct optogenetic stimulation of the injected V1 area generated significantly stronger CBV responses, with a mean active volume of 37% (range: 0.5 to 75.9%). The percentage mean active injected V1 area, over all animals, was higher for optogenetic cortical stimulation than for direct eye stimulation (optogenetic, mean: 16.2 ± 17.8 %, Wilcoxon signed-rank test, one-tailed, *p*=0.001, Fig. S1). Neurovascular activity and ChrimsonR expression were distributed similarly along the AP axis and their amplitudes were correlated (Fig. 1d, Fig. S1). Again, our control experiments demonstrated that direct optogenetic stimulation of a non-transfected cortical area resulted in no activation. The findings for these control conditions indicate that the parameters we used for optogenetic stimulation at the cortical surface did not generate CBV variations in areas into which the virus was not injected (mean: 0 ± 0%). These observations indicate that the optogenetic light stimulation used here does not trigger a vascular response detectable by fUS imaging. Consequently, with fUS imaging, we were able to visualize the entire volume of an optogenetically evoked response resulting from focal stimulation within the primary visual cortex.

We then sought to confirm that the observed changes in blood volume following optogenetic stimulation at the surface of V1 were indeed due to an increase in neuronal activity, and not to indirect factors, such as heating. We therefore performed electrophysiological recordings of V1 during the stimulation of the contralateral eye with white light (200 ms, 1 Hz, 100 cycles, 58 mW.cm^−2^) or of the injected V1 area with light at 595 nm (200 ms, 1 Hz, 100 cycles, 140 mW.mm^−2^). We used the Spyking Circus algorithm^32^ to sort the multi-unit recordings, to obtain single-cell responses. In total, we recorded a population of 171 neurons from nine animals expressing ChrimsonR in V1. These neurons displayed several distinctive patterns of activity under both direct eye and optogenetic stimulation conditions (Fig. 1e). We plotted the spike density function (SDF) of four V1 neurons for both direct eye stimulation (black lines) and optogenetic conditions (red lines), to highlight the diversity of these activity patterns. Based on the profiles of visual and optogenetic responses, we classified neurons into four different groups: neurons responding to both visual and optogenetic stimulation (V + O neurons, *n*=13, 7.6%, Wilcoxon signed-rank test, two-tailed, *p*<0.01, between baseline: [-100 0] ms, and stimulus presentation window: [0 200] ms), to visual stimulation only (V neurons, *n*=20, 11.7%), to optogenetic stimulation only (O neurons, *n*=81, 47.4%), and non-responsive neurons (‘None’ neurons, *n*=57, 33.3%). Most of the neurons responding to visual stimulation displayed phasic responses, with an ON response occurring after the start of stimulation followed, in some cases, by an OFF response. By contrast, the neurons responding to optogenetic stimulation displayed a unique ON response. We characterized the neuronal activation further, by comparing the onset latencies and durations of the V1 responses for both direct eye and optogenetic stimulation conditions, for V, O and V+O neurons (*n*=109, Fig. 1f). As observed for the representative neurons shown in Fig. 2a, the onset latencies of V1 responses were significantly shorter after optogenetic stimulation (mean: 1.76 ± 3.14 ms, *n*=76 units) than after stimulation of the contralateral eye (mean: 41.24 ± 16.61 ms, *n*=33 units, Mann-Whitney, *p*<0.0001). These very short response latencies for optogenetic stimulation are consistent with the direct activation of transfected neuronal cell bodies, bypassing all retinal synapses, by contrast to natural visual signal transmission. We also analyzed the duration of visual/optogenetic responses, to determine whether V1 neurons presented transient or sustained activity, according to the type of stimulation. For direct eye stimulation, the duration of neuronal responses was tuned on a single ensemble, with a mean duration of activation of 28.09 ± 17.05 ms (*n*=33). By contrast, in optogenetic conditions, two subgroups emerged: neurons displaying transient and sustained responses. Based on these results, we defined two subsets of neurons: transient (durations < 51 ms, *n*=33) and sustained (durations > 197 ms, *n*=22) neurons. The remaining neurons (*n*=17) had intermediate response durations. The transient responses may result from inhibitory cortical feedback from interneurons or a difference in voltage-gated channel properties between neuronal subtypes leading to the silencing of these neurons. We found no significant difference in the depth distribution of transient and sustained neurons (Fig. 1g), revealing an absence of correlation between neuronal response patterns and potential decreases in stimulus intensity with tissue depth; transient and sustained neurons were recorded within the same cortical layers (L2/3 to L6), suggesting a direct activation of cortical neurons by optogenetic stimulation at the cortical surface. We then assessed the specificity of optogenetic activation, by performing electrophysiological recordings on naive animals. In the animals in which no injection was performed (Fig. S1, *n*=3 animals), almost all the recorded neurons (*n*=187/188) displayed a total absence of response to optogenetic stimulation; the only responsive neuron had a very low response rate (3.89 Hz) relative to its baseline activity (1.71 Hz, *p* = 0.0014). In this experiment, most neurons displayed a visual response when the contralateral eye was stimulated (*n*=80/118), whereas another group of neurons (*n*=37/118) did not respond to either visual or optogenetic stimulation. Thus, by recording single-cell activities in transfected areas of V1, we were able to demonstrate that optogenetic light stimulation at the surface of the cortex triggered both an increase in cerebral blood volume, as shown by fUS imaging, and direct neuronal activation. We can therefore conclude that the fUS variations we observed reflected the optogenetic activation of V1 neurons. As in fUS imaging, we found that a larger number of neurons responded to optogenetic stimulation than to visual stimulation (Fig. 1e), consistent with the broader activation of areas in response to optogenetic stimulation than following direct contralateral eye stimulation and fUS imaging.

**Figure 2.**
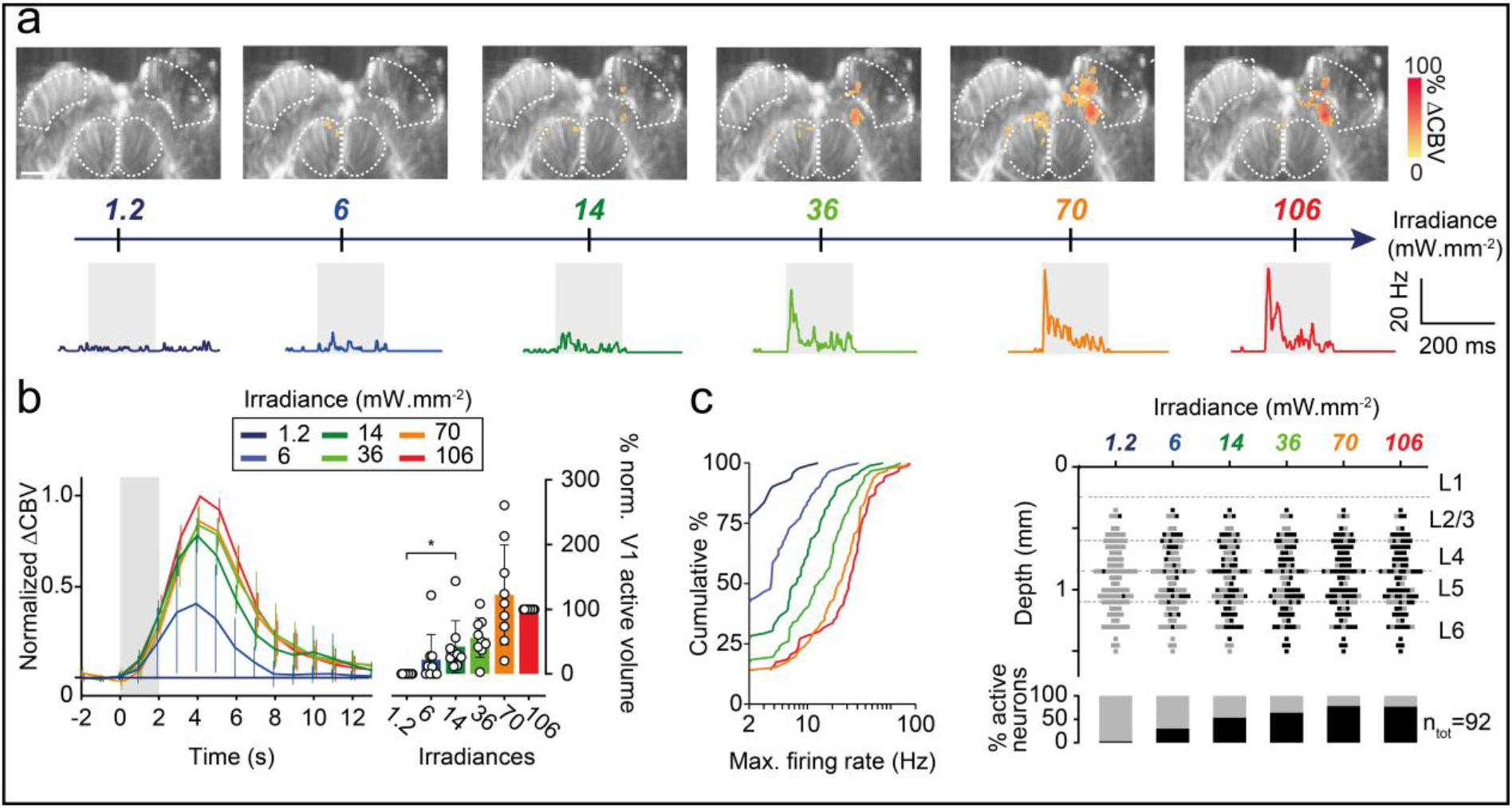
Neuronal and neurovascular optogenetic sensitivity. (a) Top: fUS activation maps of a single imaging plane (AP −7.5 mm) from animal #4 at different irradiances. Scale bar: 2 mm. White dashed lines delimit the V1 and SC areas. Bottom: SDF from a typical V1 single unit during optogenetic stimulation for 200 ms (gray patch) at different irradiances. (b) Left panel: mean CBV variation over all animals (*n*=9) for each irradiance. of the values for each animal were normalized against those obtained at 106 mW.mm-^2^. Error bars represent the standard deviation. The colored patch corresponds to the 2 s optogenetic stimulation. Right panel: normalized active volumes of the ipsilateral V1 area for each irradiance. Open circles are individual values, bars represent the mean and the standard deviation (*n*=9 animals, Wilcoxon signed-rank test, one-tailed, *p*<0.05). (c) Left: cumulative distribution of the mean maximal firing rates of all recorded V1 single units (*n*=92) during optogenetic stimulation, for each irradiance. Right panel, top: depth profile of V1 single units activated (black squares) or not (gray squares) by the optogenetic stimulation, for each irradiance (*n*=92 single units). Bottom: percentage of active (black) and non-responsive (gray) neurons for each set of conditions.

### Neuronal and neurovascular optogenetic sensitivity

As both single-cell recording and fUS imaging can be used to monitor optogenetic neuronal activation, we decided to assess the sensitivities of these two approaches. We first imaged the same single V1 plane (AP −7.5 mm) by fUS while optogenetically stimulating the surface of the ChrimsonR-expressing V1 area with various irradiances (from 1.2 to 106 mW.mm^−2^). The size of the active volume within the injected V1 area increased with irradiance, with a major increase between 36 mW.mm^−2^ and 70 mW.mm^−2^ and a slight decrease at 106 mW.mm^−2^ (Fig. 2a). One also found that the contralateral SC was slightly activated at irradiances above 1.2 mW.mm^−2^. For each irradiance, we then calculated the mean CBV variation over all significant pixels in the V1 area and over all trials (Fig. 2b, left panel). For each animal, we normalized these values against those obtained for the highest irradiance (106 mW.mm^−2)^. Normalized CBV variation peaked 4 s after the start of stimulation. Peak CBV values increased as a function of irradiance. No CBV variation was recorded for the lowest irradiance (1.2 mW.mm^−2^). Difference in peak CBV values relative to that for the lowest irradiance tested started to become significant from 6 mW.mm^−2^ onwards (1.2 mW.mm^−2^, mean: 0 ± 0; 6 mW.mm^−2^, mean: 0.36 ± 0.34, Wilcoxon signed-rank test, one-tailed, *p*<0.05). For each irradiance, we then calculated the percentage normalized activated ipsilateral V1 volume (Fig. 2b, right panel) in all animals (*n*=9). We again observed an increase in the normalized active volume of V1 with irradiance. Difference with respect the value obtained at the lowest irradiance became significant from 6 mW.mm^−2^ onwards (1.2 mW.mm^−2^, mean: 0 ± 0%; 6 mW.mm^−2^, mean: 22.3 ± 39.0%, Wilcoxon signed-rank test, one-tailed, *p*<0.05). We then investigated whether the variation of CBV responses observed with irradiance was related to the number of neurons recruited. The SDF from a single-unit example was determined for the different irradiances; the amplitude of the neuronal response increased with irradiance from 8.2 ± Hz for 6 mW.mm^−2^ to 36.6 ± 73 Hz for 106 mW.mm^−2^ (Fig. 2a). We generalized this analysis by plotting the cumulative distribution of the mean maximal firing rates for each irradiance (Fig. 2c, left panel). The cumulative curves reached a plateau for lower maximal firing rates with decreasing irradiance. We noted a significant difference in distribution between 1.2 mW.mm^−2^ and irradiances of 6 mW.mm^−2^ and above (1.2 mW.mm^−2^, median: 0 Hz; 6 mW.mm^−2^, 3.8 Hz, Kolmogorov-Smirnov test, *p*<0.0001), demonstrating that electrophysiological recordings and fUS imaging had equivalent sensitivities. Finally, in our total population of V1 neurons, the proportion of responsive neurons increased with irradiance (from 28/92 units at 6 mW.mm^−2^ to 71/92 units at 106 mW.mm^−2^, Fig. 2c, right panel). Interestingly, the depth distribution of the activated neurons did not change with increasing irradiance (Fig. 2c, right panel). It was, therefore, possible to activate neurons from deep cortical layers even at very low irradiances.

### Spread of optogenetic activation to downstream and upstream visual areas

The results described above relate to optogenetic activation in V1 with the optic fiber placed at the surface of the primary visual cortex. We then investigated whether the activity initiated in V1 spread to other visual structures up- or downstream. This aspect is important for optogenetic therapies for the restoration of cortical vision because it would demonstrate the propagation of visual information favoring the generation of visual percepts. For the downstream pathway, LGN terminals end in V1 at the depth of cortical layer IV, whereas cortical V1 layer VI sends feedback connections to the LGN. We thus hypothesized that injecting AAV9-7m8-CaMKII-ChrimsonR-tdT into V1 would increase ChrimsonR expression in at least one of these two categories of fibers and that the optogenetic activation of V1 would lead to direct activation of the LGN via LGN terminals in V1, or to an indirect activation of the LGN through feedback connections in V1 cortical layer VI. Histological analyses confirmed that some ChrimsonR expression occurred in the LGN, but we were unable to identify any ChrimsonR-expressing cell bodies in the LGN, suggesting that only retrograde fibers originating from cortical V1 layer VI expressed this opsin (Fig. S2). We imaged the planes containing the LGN (AP −3.5 to AP −5.5 mm). Figure 3a shows the fUS imaging planes at AP −5 mm for the direct eye and optogenetic stimulation conditions. We noted a slight activation of the visual cortex following visual stimulation (see Fig. 1b), and a strong activation of the ipsilateral LGN (*n*=85/225 active pixels) when the contralateral eye was stimulated with the white LED, suggesting that this kind of stimulation was more appropriate for LGN and SC activations than for the visual cortex. Indeed, in the 11 animals (Fig. 3b), the mean percentage of the LGN volume visually activated was 20.5 ± 13.7% which is much greater than the volume activation obtained for V1 (less than 1%). When the injected V1 surface was stimulated with the optic fiber (Fig. 3a), CBV responses increased significantly in the ipsilateral LGN, but with a much smaller number of activated pixels (*n*=12/225 pixels) than for visual stimulation. We also performed a control optogenetic stimulation, in which we stimulated the non-injected V1 area. We observed no activation of the ipsilateral LGN, confirming that, on stimulation of the injected hemisphere, ipsilateral LGN activation was due to optogenetic activation of LGN terminals in V1 or feedback from the V1 area (Fig. S2). In the 11 animals tested, the active LGN volumes for visual stimulation were larger than those for optogenetic stimulation (visual, mean: 20.5 ± 13.7%; optogenetic, mean: 6.5 ± 12.3%, Wilcoxon signed-rank test, *p*=0.0068). In addition, for the five animals tested by optogenetic stimulation of the non-transfected V1 area, we detected no significant responsive pixels (Fig. 3b, mean: 0 ± 0%, *n*=5 animals). We also performed single-cell recordings in the LGN, to demonstrate that the neurovascular activations imaged by fUS in the LGN coincided with the activation of LGN neurons (Fig. 3c). We recorded a total of 153 neurons in the LGN. Only two units responded to both visual and optogenetic stimulation; seven units were responsive only to optogenetic stimulation, and 40 units were responsive only to visual stimulation (*n*=7 animals). The vast majority of LGN neurons were unresponsive to both types of stimulation. The distribution of onset latencies following direct eye stimulation was broader for LGN than for V1 single units (Fig. 3d, mean: 47.88 ± 23.55 ms, *n*=42 units, and see Fig. 1f). This may reflect the recording of both cells activated by retinal ganglion cells and cells retrogradely activated by the V1 area, which have higher latencies. Following optogenetic stimulation at the surface of the V1 area, the onset latencies for LGN neurons were shorter than those following visual stimulation (mean: 9.86 ± 4.95 ms, *n*=7 units, Mann-Whitney, *p*<0.05). However, these response latencies were greater than those recorded in V1 (*n*=42 units. Mann-Whitney, *p*<0.0001, see Fig 1f for V1 units). This result suggests that the optogenetically activated LGN single units recorded here were activated indirectly, by retrograde fibers from V1 cortical layer VI, as suggested by the histological data.

**Figure 3.**
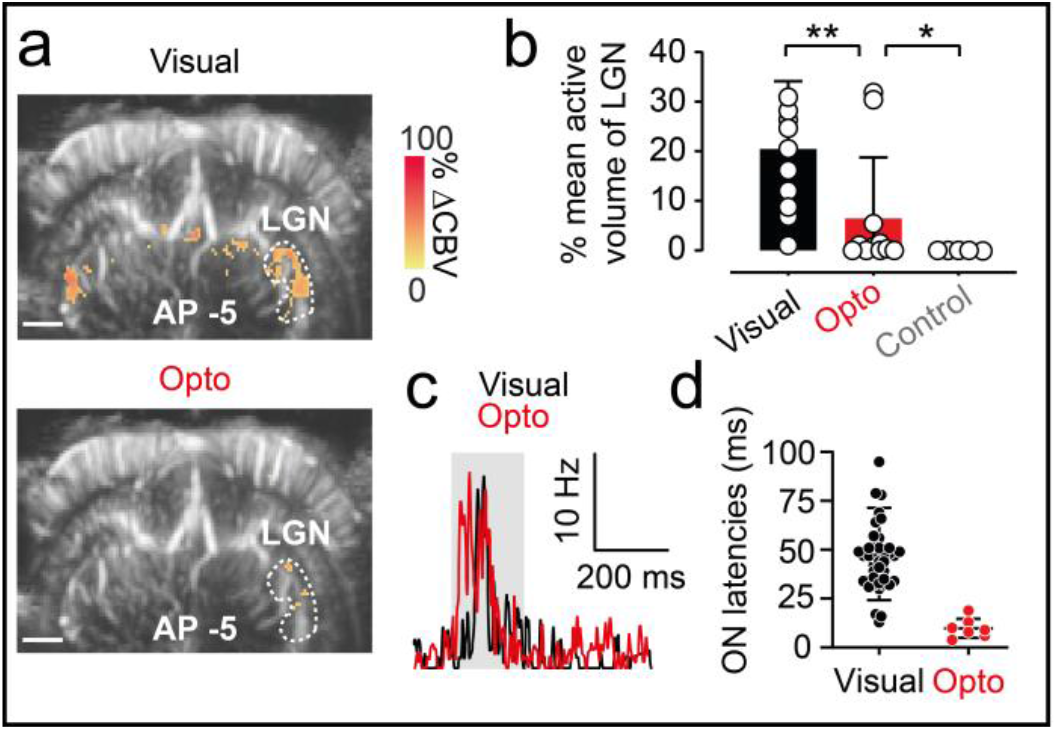
Spread of optogenetic activation to the LGN. (a) fUS activation maps obtained during visual stimulation of the contralateral eye (top) and optogenetic stimulation of the ChrimsonR-expressing ipsilateral V1 area (bottom) from a single imaging plane in which the LGN is present (AP −5 mm) from the same animal. White dashed lines delimit the ipsilateral LGN. Scale bar: 2 mm. (b) Percentage active volume of the LGN after visual, optogenetic or control stimulation. For both visual and optogenetic stimulations, we show the volumes of the LGN ipsilateral to the injection, whereas, for control stimulations, the volume of the contralateral LGN is shown. Open circles represent mean values over all imaging planes for each animal (visual and optogenetic stimulation, *n*=11, Wilcoxon signed-rank test, one-tailed, *p*=0.0068, *n*=5 for control, Mann-Whitney test, one-tailed, *p*=0.0288), bars represent the mean for all animals. (c) SDF of a typical LGN single unit responding to visual (black line) and optogenetic (red line) stimulation of the ipsilateral V1 area. (d) ON latencies of V1 and LGN single-unit responses to visual (V1, *n*=33 units, LGN, *n*=42 units) or optogenetic stimulation of V1 (V1, *n*=29 units, LGN, *n*=7 units. Mann-Whitney test, *p*<0.05). The irradiance used for optogenetic and control stimulation was 142 mW.mm^−2^.

We also investigated whether V1 optogenetic stimulation could spread to the direct upstream visual area toward which V1 projects: the V2 area. We obtained fUS activation maps in different imaging planes (AP −6 to AP −8 mm) in which the entire V2 area was present after either direct stimulation of the contralateral eye or optogenetic stimulation at the surface of the V1 area. We show representative activation maps from single imaging planes (AP −8.5 mm) in Figure 4a. Visual stimulation of the contralateral eye led to almost no activation of the V2 area on the same side as the injected V1. By contrast, optogenetic stimulation of the transfected V1 area led to a stronger activation of the ipsilateral V2, mostly within the ventral part of V2. As previously reported, within the different imaging planes, we also observed strong activation in the lower parts of V1 and V2 containing the axons. We next quantified the active volume over all imaging planes for the V2 area, for all 10 animals (Fig. 4b). As for the V1 area, visual stimulation resulted in only weak activation of V2 (mean: 0.2 ± 0.3% of activated volume). Averaged activation volumes were significantly larger for optogenetic stimulation than for visual stimulation (mean: 5.6 ± 8.0% activated volume, *n*=10 animals, Wilcoxon signed-rank test, one-tailed, *p*<0.05). Control stimulation of the non-injected V1 area confirmed the specificity of the spread of optogenetic activation from V1 to V2, as no activation was observed in the contralateral V2 area (Fig. 4b, Fig. S2).

**Figure 4.**
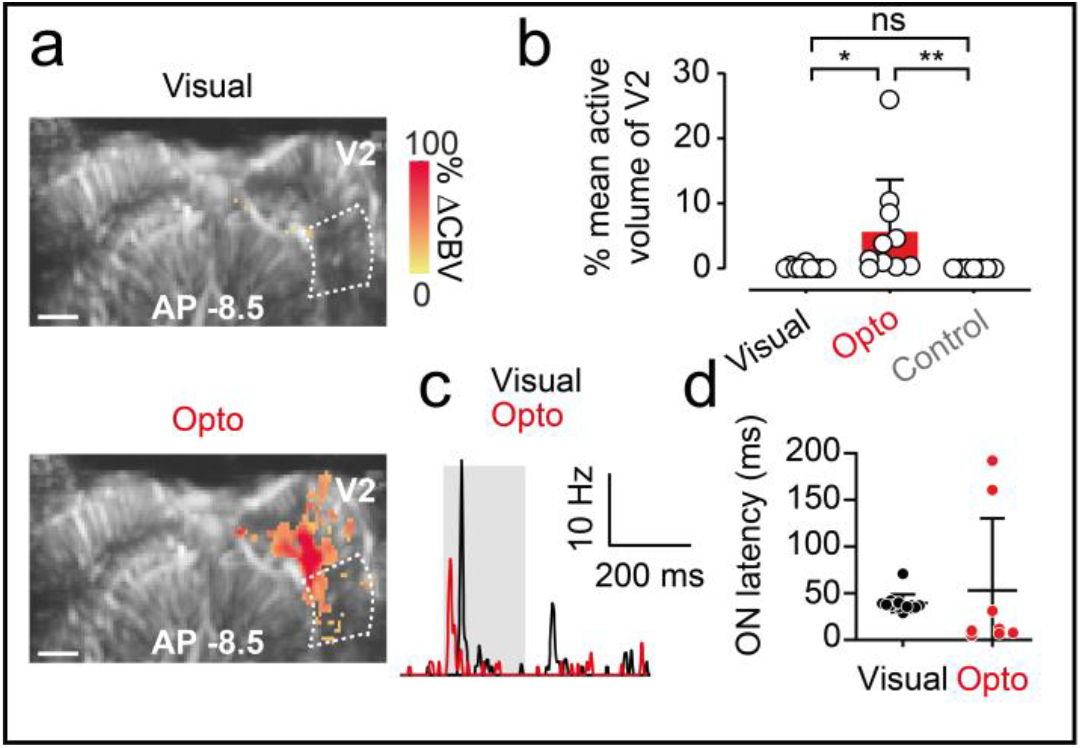
Spread of the optogenetic activation to V2. (a) fUS activation maps obtained during visual stimulation of the contralateral eye (top) and optogenetic stimulation of the ChrimsonR-expressing ipsilateral V1 area (bottom) and control stimulation of the uninjected contralateral area from a single imaging plane (AP −7.5 mm) containing the V2 area. White dashed lines delimit the ipsilateral V2 area. Scale bar: 2 mm. (b) Percentage active volume of V2 after visual, optogenetic or control stimulation. For both visual and optogenetic stimulation, we show the volumes of V2 ipsilateral to the injection, whereas, for the control, we shown the volume of the contralateral V2. Open circles represent mean values over all imaging planes for each animal (visual and optogenetic, *n*=1, Wilcoxon signed-rank test, one-tailed, *p*=0.002, *n*=7 for control, Mann-Whitney test, one-tailed, *p*=0.0004), bars represent the mean over all animals. (c) SDF of a typical V2 single unit responding to visual (black line) and optogenetic (red line) stimulation of the ipsilateral V1 area. (d) ON latencies of V1 and V2 single-unit responses to visual (V1, *n*=33 units, V2, *n*=14 units) or optogenetic stimulation of V1 (V1, *n*=76 units, V2, *n*=8 units). The irradiance used for optogenetic and control stimulation was 140 mW.mm^−2^.

Electrophysiological recordings within V2 confirmed that the fUS variations we observed in V2 were consistent with direct neuronal activation (Fig. 4c). We recorded V2 single units responding to visual and optogenetic stimulation. As previously described, we characterized the onset latency of these neurons, comparing the values obtained with those for V1 single units (Fig. 4d). Similar visual latencies were recorded for V1 and V2 single units (V1, mean: 41.24 ± 16.61 ms, *n*=33 units (see Fig. 1f); V2, mean: 39.43 ± 9.557 ms, *n*=14 units). Interestingly, optogenetic onset latencies were higher for V2 single units than for V1 (V1, mean: 1.76 ± 3.14 ms, *n*=76 units. V2, mean: 62.78 ± 77.44 ms, *n*=9 units, Mann-Whitney, *p*<0.0001), suggesting that these neurons were activated indirectly by ChrimsonR-expressing V1 neurons. Two V2 neurons presented particularly long optogenetic ON latencies, possibly due to a difference in the microcircuits involved. We checked that this variability did not bias the latency delay for V2 and V1 neurons after optogenetic stimulation, by performing the same statistical test on a V2 data set restricted to the remaining seven fast V2 units. The difference in ON latency between V1 neurons and these fast V2 neurons was conserved (fast V2 units, mean: 12.2 ± 9.5 ms, *n*=7 units, Mann-Whitney, *p*<0.0001), suggesting that all V2 neurons were activated indirectly after the onset of stimulation. This conclusion is supported by the lack of ChrimsonR expression in V2 on brain slices from the animals used to record these single units. However, these V2 latencies were shorter than V2 latencies for natural eye stimulation, demonstrating that they were genuinely produced by V1 optogenetic activation.

## Discussion

### Coupling of optogenetics with fUS

In this study, we demonstrate that optogenetic activation can be detected by fUS imaging. Rungta and colleagues^30^ indicated that blue light from the tip of the optic fiber *per se* could induce neurovascular responses in naïve mice, at irradiances higher than 2 mW ∼ 18 mW.mm^−2^. Another recent study reported hemodynamic responses following blue light illumination of the retrosplenial cortex in Thy-ChR2 mice or the illumination of specific neuronal subpopulations of the superior colliculus in other transgenic lines^26^. The authors explained that they used a lower light power and irradiance (0.3 mW ∼1.5 mW.mm^−2^) to prevent non-specific activation. By contrast, we used an AAV-mediated strategy to express ChrimsonR in V1 neurons, resulting in a lower density of opsin-expressing cells than in transgenic mice lines^33^. This generalizes the possibility of recording neurovascular optogenetic activation due to a small number of transfected neurons. Moreover, we show here that stimulation of the control non-injected hemisphere does not induce a vascular response, thereby demonstrating the specificity of ChrimsonR-elicited optogenetic activity. Single-cell recordings confirmed the neuronal optogenetic activation correlated with these CBV variations. We also used a higher wavelength (595 nm) for illumination than Rungta and coworkers, but with comparable values of power and irradiance. These results indicate that red light can be used even at high power, in protocols combining the optogenetic activation of red-shifted opsins and fUS imaging. Importantly, electrophysiological recordings from the control animals showed no neuronal activation, suggesting negligible thermal effects with the use of 595 nm light under the parameters used here. Indeed, we included these parameters in the heat propagation model of Stujenske^34^, and recorded a very local (<1 mm-diameter sphere from the tip of the optic fiber) increase in temperature, by 0.2°C, which is too small to alter neuronal firing rates. Finally, we characterized the dose-response dependence of our recorded optogenetic activations. We obtained equivalent sensitivities for optogenetic responses measured by fUS imaging and by electrophysiological recordings.

The use of fUS imaging to analyze optogenetic responses has the advantage that it could potentially provide a mesoscopic view of the activated area. Unlike electrophysiological recordings, which extract information very locally^20^, and calcium imaging, which is dependent on both expression of the calcium indicator and the penetration of blue light through the tissue^35^, it can display information at a brain-wide spatial scale. The coupling of fMRI with optogenetics meets these criteria, but with a lower spatial resolution^21,22,36^ (of the order of a millimeter per pixel). By contrast, fUS imaging can detect brain activity at a submesoscopic resolution (100 x 100 µm^2^), with less cumbersome equipment than for fMRI.

### Spread of the activity

We found that optogenetic neurovascular activation was well correlated with ChrimsonR expression in the AP axis. We also noted that optogenetic V1 stimulation spread beyond the borders of V1, as indicated by the activation of both the LGN and V2 on both fUS and electrophysiology. This spread of activity may be due to fibers located at the base of V1, connected to the LGN and expressing ChrimsonR (Fig. S1). These fibers may generate the neurovascular activity detected ventrally to V1 on our activation maps. Furthermore, the mean neuronal transfection rate was 5.5% in the coronal plane displaying the highest fluorescence signal, whereas a larger volume of V1 was activated in this plane. A first explanation for this may be the spread of neuronal activation to other V1 cells, although we recorded a rather homogeneous set of short-onset latencies. Alternatively, neurovascular activations may be broader *per se* than neuronal responses. Some studies of rat olfactory bulb glomeruli have provided evidence of a close overlap between capillary blood flow and neuronal activity^37^, whereas others have reported a mismatch between the areas of neurovascular and neuronal activation^38,39^. Specifically for V1, a lack of correlation between BOLD signals and spiking activity has been observed in cats^40^, and a lack of correlation between single-vessel hemodynamic responses and calcium imaging signals has been found in cats and rats^41^. Importantly, we show here that the neurovascular and neuronal activities initiated in V1 spread to the LGN and V2. This propagation of visual information is important for optogenetic cortical vision restoration therapy, because it favors the generation of visual percepts.

### Cortical visual restoration

We detected neurovascular and neuronal responses, even at low irradiance (6 mW.mm^−2^), with no modification of the depth distribution of the activated neurons. One recent study^42^ reported that the stimulation of deep V1 layers (>1.5 mm) in non-human primates with electrical prostheses elicited behavioral responses. Our ability to activate neurons in deep cortical layers highlights the potential value of red-shifted opsin ChrimsonR for optogenetic cortical vision restoration strategies. A key element of visual restoration is the induction of neuronal responses with characteristics matching those resulting from a natural visual stimulus^5–11^. We show here that the firing rates induced by optogenetic stimulation were similar to those induced by visual stimulation. Optogenetically activated neurons had very short latencies, of the order of 1-2 ms. Theoretically, this is sufficient for the encoding of natural images into optogenetic stimulations at a temporal resolution matching the resolution of the natural visual signal. Moreover, those optogenetic onset latencies were quite homogeneous, despite being obtained for neurons located in different layers. These results demonstrate that we can induce a signal that does not lose its temporal resolution with depth.

Finally, the feasibility of inducing visual percepts by optogenetic cortical vision restoration remains to be demonstrated. In species with a more complex hierarchical organization of cortical visual areas, such as nonhuman primates, a few studies have shown behavioral effects due to the optogenetic stimulation of higher visual areas. Jazayeri^43^ and coworkers reported that saccades following fixation tasks were shifted towards the receptive field of the region of V1 optogenetically activated following the fixation point offset. Ju^44^ and coworkers demonstrated the successful detection of optogenetic percepts in a saccade task. Our ability to detect a spread of activity from V1 to other visual areas is consistent with these behavioral studies indicating perception. Here, we used a single optic fiber for optogenetic stimulation. Replacing this device with a more complex stimulation system, such as arrays of micro-LEDs, might make it possible to generate percepts more complex than phosphenes and to develop discrimination behavioral tasks to assess their perception.

## Materials and methods

### Animals

All animal experiments and procedures were approved by the Local Animal Ethics Committee (registration number 2018032911282465) and performed in accordance with European Directive 2010/63/UE. We used wild-type male Long-Evans rats (Janvier Laboratories), nine weeks old at the time of viral injection. Rats were maintained under a reverse 12-hour light/12-hour dark cycle, with ad libitum access to food and water, except during surgery and electrophysiological recordings.

### AAV production

The AAV9-7m8-CaMKII-ChrimsonR-tdT vector was packaged as previously described, by the triple transfection method, and purified by iodixanol gradient ultracentrifugation^45^. The AAV9-7m8-CaMKII-ChrimsonR-tdT vector was titered by qPCR with SYBR Green^46^ (Thermo Fisher Scientific). The titer used in this study was 4.39 x 10^12^ vg.mL^−1^.

### Immunostaining and confocal imaging

Following electrophysiological recordings, rats were euthanized, and their brains were extracted and fixed by overnight incubation in 4% paraformaldehyde (100496, Sigma-Aldrich) at 4°C. Brains were then cryoprotected in 30% sucrose (84097, Sigma-Aldrich) and 50 µm sagittal slices were cut with a microtome (HM450, Microm). The slices with the most intense tdT fluorescence from each brain were selected for further immunohistochemistry and imaging. Cryosections were permeabilized by incubation with 0.5% Triton X-100 in PBS for 1 h at room temperature and were then incubated in blocking buffer (PBS + 1% BSA + 0.1% Tween 20) for 1 hour at room temperature. Samples were incubated overnight at 4°C with monoclonal anti-NeuN antibody (1:500; Mouse, MAB377, Merck Millipore), in a 50% dilution of blocking buffer + 0.5% Triton X-100. Secondary antibodies conjugated with Alexa Fluor dyes (1:500; Molecular Probes) and DAPI (1:1000, D9542, Merck Millipore), were incubated with the samples for 1 hour at room temperature. An Olympus FV1000 laser-scanning confocal microscope with a 20x or 40x objective (UPLSAPO 20XO, NA: 0.85) was used to acquire images of brain sections.

### Viral injections

Viral injections were performed in aseptic conditions with a digital small-animal stereotaxic instrument (David Kopf Instruments). Ear bars were covered with xylocaine to ensure that the animals felt no pain. Rats were anesthetized in a sealed box containing gaseous isoflurane (5%), and maintained under anesthesia in the stereotaxic frame (25% ketamine, 10% medetomidine and 65% saline injected intraperitoneally) for the entire surgical procedure, and animal body temperature was maintained at 37°C with a heating pad. Buprenorphine was injected subcutaneously to reduce inflammation, and Lubrithal was applied to the eyes to prevent them from drying out. Xylocaine, 70% ethanol and Vetedine were applied successively to the scalp before incision, to minimize pain and maintain sterile conditions. Cranial sutures were cleaned to remove connective tissue, by applying H_2_O_2_, to facilitate localization of the injection coordinates. We injected a total volume of 1.2 µL of viral suspension unilaterally into rates, via two injection tracks, at a flow rate of 50-75 nL/min, with a 5 or 10 µL microsyringe (Hamilton) equipped with a microinjector (Sutter Instrument) and controller (World Precision Instruments). The coordinates for viral injection were +2.8 / +3.2 mm from midline (M-L axis), −6.5 / −7.5 mm from Bregma (A-P axis) and 1.6-1.35-1.1 mm ventral to the skull surface (D-V axis), based on the 2004 edition of the Paxinos and Franklin rat brain atlas. Viral efflux was prevented by leaving the needle in a 1.8 mm ventral position for two minutes before beginning the injection, with a three-minute interval left between injections before complete withdrawal of the needle from the cortex. After surgery, rats were brought round from anesthesia with a subcutaneous injection of Antisédan (0.15 mL).

### *In vivo* electrophysiological recordings

Bilateral craniotomies were performed with a digital small-animal stereotaxic instrument (David Kopf Instruments), at least 30 days after viral injection, to allow time for opsin expression. Ear bars were covered with xylocaine to prevent pain. Rats were anesthetized in a sealed box containing gaseous isoflurane (5%) and maintained under anesthesia in the stereotaxic frame (25% ketamine, 10% medetomidine and 65% saline injected intraperitoneally) for the entire surgical procedure, and electrophysiological recordings were taken with body temperature maintained at 37°C with a heating pad. Buprenorphine was injected subcutaneously to reduce inflammation, and Lubrithal was applied to the eyes to prevent them from drying out. Cranial sutures were cleaned to remove connective tissue, by applying H_2_O_2_, to reveal the injection tracks. Parietal bones were removed by drilling rectangular flaps and gently moving the bone away from the dura mater, exposing the cortex from 3 to 8.5 mm from Bregma (A-P axis), to cover the injection tracks. The dura was then gently removed. During drilling, the skull was regularly cooled with PBS, and once the cortex was exposed, it was protected from dehydration by the regular application of cortex buffer. After surgery, electrophysiological recordings were performed with 16-channel electrodes (A1×16-5 mm-50-703-OA16LP) coupled with a 400 µm-core fiber (Thorlabs M79L005 Fiber Cable, MM, 400 µm 0.39NA, FC/PC to 1.25 mm ferrule, 0.5 m) at various positions close to the injection sites. Electrophysiological data were acquired with MC RACK software. For visual stimulation, a white LED (Thorlabs MNWHL4 Mounted LED, 5V, 60 mW/cm^2^) was placed 15 cm in front of the eye on the contralateral side of the cranial window.

### Acquisition protocol for electrophysiological recordings

For optogenetic stimulation, we connected the optic fiber (reference above) to a light source (Thorlabs M595F2 (fiber coupled LED @595 nm), Ø400 µm, 150 µW/cm^2^) delivering light at the ChrimsonR excitation wavelength (595 nm). We targeted the transfected region of V1 by imaging tdT fluorescence with a Micron IV imaging microscope (Phoenix Research Laboratories) before recordings. The fiber was placed on the surface of the cortex while the electrode was inserted in the tissue. Both stimulations consisted of 100 repeats of 200 ms flashes at 1 Hz. The onset of the flashes was aligned with electrophysiological data, with Clampex 9.2 software. We used several different irradiances of light at 595 nm in this study. Power at the fiber tip was measured with a power meter (Thorlabs, PM100D), by placing the fiber tip in contact with the sensor. The irradiance corresponding to each power was calculated as previously described^30^.

**Table 1:** 595 nm light irradiances and powers used in this study.

| Irradiance (mW.mm<sup>-2</sup>) | Power (mW) |
| --- | --- |
| 142 | 7.1 |
| 140 | 7.0 |
| 106 | 5.3 |
| 70 | 3.5 |
| 36 | 1.8 |
| 14 | 0.7 |
| 6 | 0.3 |
| 1.2 | 0.06 |

### Spike sorting

Offline spike sorting of the electrophysiological recordings (linear 16-channel electrodes) was performed with the SpyKING CIRCUS package^32^. Raw data were first high-pass filtered (> 300 Hz) and spikes were detected when a filtered voltage trace crossed the threshold. Automatic cluster extraction was performed and candidate clusters were curated. Refractory period violations (< 2 ms, >1% violation) and noisy spike shapes led to cluster deletion. Spike templates with coordinated refractory periods in the cross-correlogram together with similar waveforms led to the merging of cluster pairs.

### Electrophysiological data analysis

All electrophysiological data were extracted and analyzed with a custom-made Matlab script. Responsive units were defined as those displaying a significant difference in neuronal activity between the pre-stimulation period (averaged 100 ms before stimulus onset) and the stimulation interval (averaged 200 ms following stimulus onset, Wilcoxon signed-rank test, *p*<0.01). For each unit, responses are represented as the spike density function (SDF), which was calculated from the mean peristimulus time histogram (PSTH, bin size: 1 ms, 100 trials) smoothed with a Gaussian filter (2 ms SD).

### Calculation of latencies and response durations

The latency of the units displaying activation was defined as the first time point at which the SDF crossed the value of the baseline plus 2SD and remained higher than this value for at least 10 ms. Conversely, the offset of activation was defined as the first time point after latency that the SDF crossed back below the value of the baseline plus 2SD and remained lower than this value for at least 10 ms. Not all active neurons from a given population met these criteria, accounting for the slight difference between the number of active neurons and the number of latencies presented here.

### Generation of functional ultrasound images

fUS imaging was performed as previously described^24^, with a linear ultrasound probe (128 elements, 15 MHz, 110 µm pitch and 8 mm elevation focus, Vermon; Tour, France) driven by an ultrafast ultrasound scanner (Aixplorer, Supersonic Imagine; Aix-en-Provence, France).

### Acquisition protocol for fUS imaging

3D fUS acquisitions were performed after craniotomy and electrophysiological recordings, as previously described. When optogenetic responses were observed, the position of the optic fiber on the surface of the cortex was kept unchanged until control acquisitions were performed, in which the fiber was moved to the other hemisphere. The cortex buffer on the surface of the cortex dried out, and 1 cm^3^ of ultrasound coupling gel was placed between the cortex and the linear ultrasound probe. Acquisition protocols consisted of 20 stimulation blocks, each consisting of 13 s of rest followed by 2 s of stimulation. For visual stimulation, the white LED used for electrophysiological recordings was kept in the same position, and the 2 s stimulation consisted of eight repeats of 125 ms flashes at 4 Hz. For optogenetic stimulation, the 2 s stimulation consisted of 40 repeats of 25 ms flashes at 20 Hz. Acquisitions were performed on coronal planes from 3 mm to 8.5 mm from Bregma (AP axis), with a 0.5 mm increment corresponding to the thickness of the imaging plane.

### Building activation maps

For each pixel, we averaged, for each block, the intensity of Doppler power at baseline (2 s before stimulus onset) and during a response window (4 s after stimulus onset). The signals were then compared in one-tailed Wilcoxon signed-rank tests. Only pixels with *p*-values < 4.03 x 10^−6^ (corresponding to a global *p*-value < 0.05 with Bonferroni correction) were considered significant. On the maps, CBV during the response window is presented as a percentage of the baseline value. Region of interest (ROI) as V1, V2, SC and LGN were determined for each imaging plane, based on the Matt Gaidica rat brain atlas.

## Acknowledgments

We thank Pierre Pouget, Frédéric Chavane, Brice Bathellier, Kévin Blaize, Sara Cadoni and Diep Nguyen for their useful discussions about the data. We thank Julie Dégardin for her help with surgery, Baptiste Lefebvre for assistance with spike sorting and Hicham Serroune for technical assistance with fUS imaging. This work was supported by the European Research Council (ERC) Synergy Grant Scheme (ERC Grant Agreement 610110).

## Author contributions

M.P., S.P., J-A.S., M.T., G.G., and F.A. designed the study; M.P. and D.D. designed viral vectors; M.P. and W.D. produced viral vectors; M.P., G.L., A.R., R.G. and M.V. performed intracortical injections; M.P., G.L. and C.J. performed electrophysiological recordings and fUS acquisitions; G.L. performed histological experiments; M.P., U.F., A.C., G.G., F.A. and S.P. analyzed data; M.P., G.G. and F.A. constructed the figures; M.P., A.C., G.G., F.A. and S.P. wrote the manuscript.

