## Supplementary figures and images for "Functional ultrasound imaging of the spreading activity following optogenetic stimulation of the rat visual cortex"

### FigS1

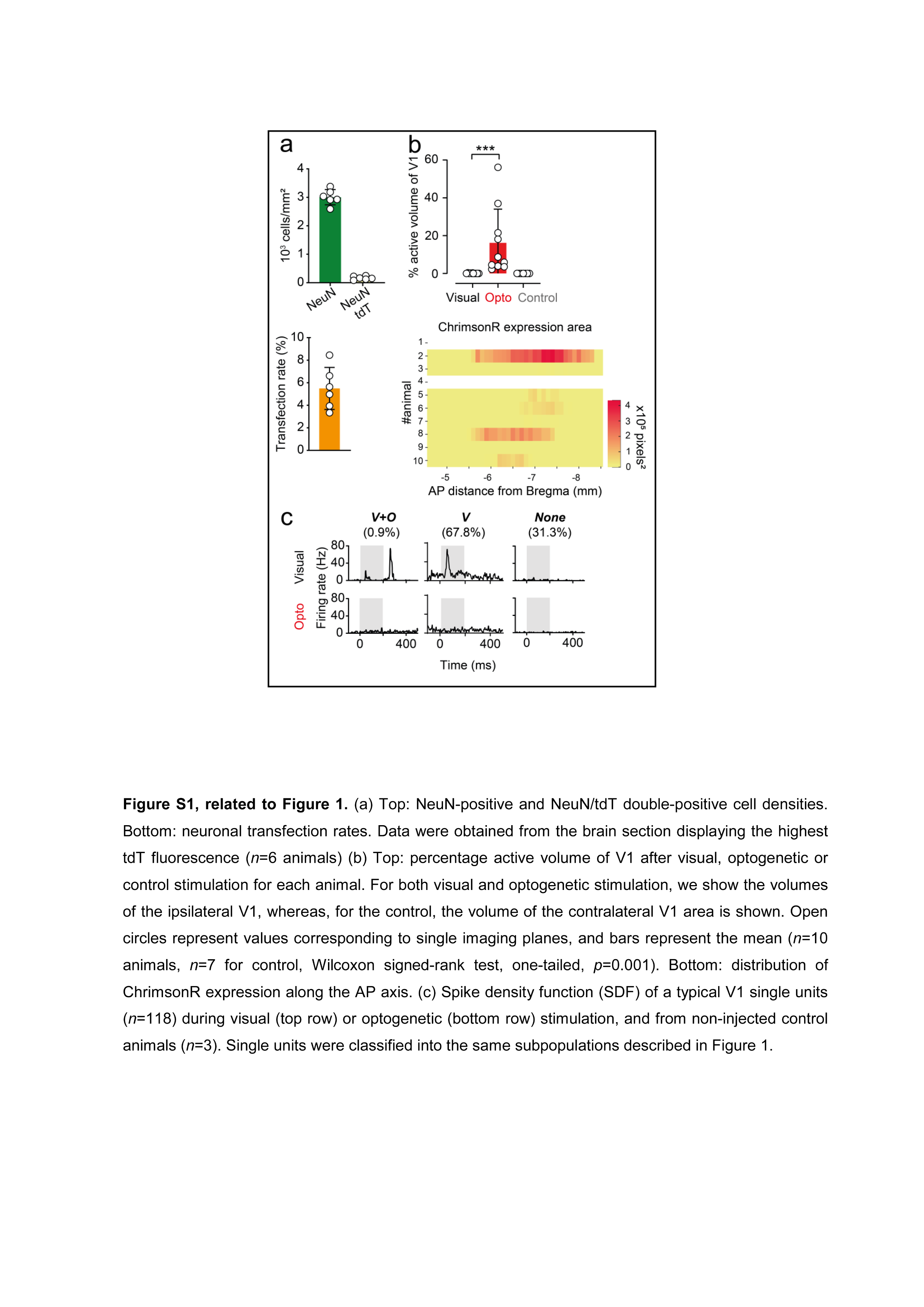

### FigS2

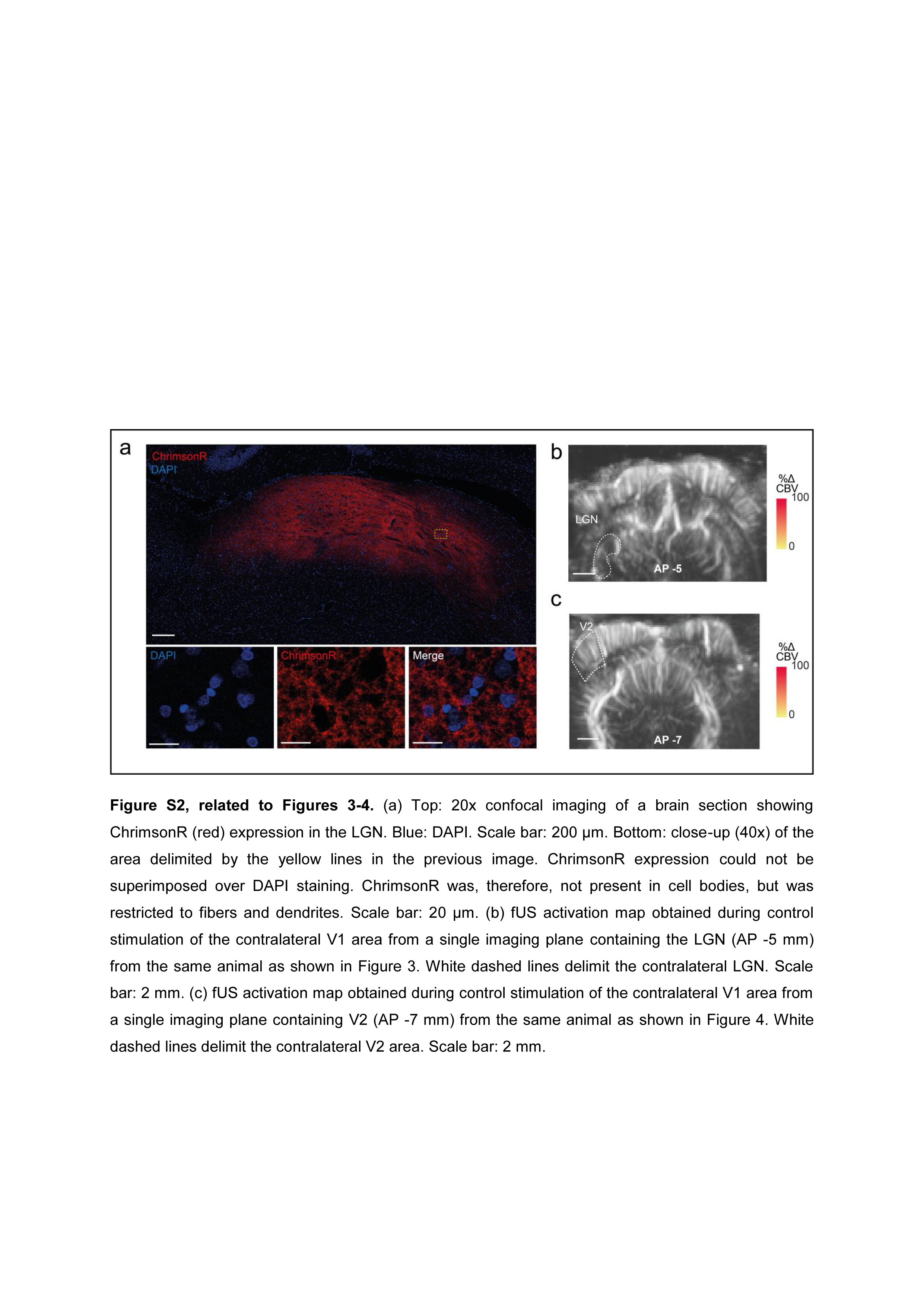
